# Quantitative assessment of fecal contamination in multiple environmental sample types in urban communities in Dhaka, Bangladesh using SaniPath microbial approach

**DOI:** 10.1101/723528

**Authors:** Nuhu Amin, Mahbubur Rahman, Suraja Raj, Shahjahan Ali, Jamie Green, Shimul Das, Solaiman Doza, Momenul Haque Mondol, Yuke Wang, Mohammad Aminul Islam, Mahbub-Ul Alam, Tarique Md. Nurul Huda, Sabrina Haque, Leanne Unicomb, George Joseph, Christine L. Moe

## Abstract

Rapid urbanization has led to a growing sanitation crisis in urban areas of Bangladesh and potential exposure to fecal contamination in the urban environment due to inadequate sanitation and poor fecal sludge management. Limited data are available on environmental fecal contamination associated with different exposure pathways in urban Dhaka. We conducted a cross-sectional study to explore the magnitude of fecal contamination in the environment in low-income, high-income, and transient/floating neighborhoods in urban Dhaka. Ten samples were collected from each of 10 environmental compartments in 10 different neighborhoods (4 low-income, 4 high-income and 2 transient/floating neighborhoods). These 1,000 samples were analyzed with the IDEXX-Quanti-Tray technique to determine most-probable-number (MPN) of *E. coli*. Samples of open drains (6.91 log_10_ MPN/100 mL), surface water (5.28 log_10_ MPN/100 mL), floodwater (4.60 log_10_ MPN/100 mL), produce (3.19 log_10_ MPN/serving), soil (2.29 log_10_ MPN/gram), and street food (1.79 log_10_ MPN/gram) had the highest mean log_10_ *E. coli* contamination compared to other samples. The contamination concentrations did not differ between low-income and high-income neighborhoods for shared latrine swabs, open drains, municipal water, produce, and street foodsamples. *E. coli* contamination were significantly higher (p <0.05) in low-income neighborhoods compared to high-income for soil (0.91 log_10_ MPN/gram, 95% CI, 0.39, 1.43), bathing water (0.98 log_10_ MPN/100 mL, 95% CI, 0.41, 1.54), non-municipal water (0.64 log_10_ MPN/100 mL, 95% CI, 0.24, 1.04), surface water (1.92 log_10_ MPN/100 mL, 95% CI, 1.44, 2.40), and floodwater (0.48 log_10_ MPN/100 mL, 95% CI, 0.03, 0.92) samples. *E. coli* contamination were significantly higher (p<0.05) in low-income neighborhoods compared to transient/floating neighborhoods for drain water, bathing water, non-municipal water and surface water. Future studies should examine behavior that brings people into contact with the environment and assess the extent of exposure to fecal contamination in the environment through multiple pathways and associated risks.

## Introduction

Globally, an estimated 24% of the total disease burden and 23% of all deaths are attributed to environmental factors [1]. Inadequate sanitation and unsafe fecal sludge management threaten public health through fecal contamination in the environment in many low- and middle-income countries [2,3]. Dhaka, the capital of Bangladesh, is one of the most densely populated cities in the world [4]. Fecal contamination in the environment is common in Dhaka neighborhoods due to many factors, including poor sanitation and sewerage systems, rapid unplanned urbanization, frequent flooding [5,6], and inefficient solid waste management [7,8]. Recent studies in urban Dhaka [9] and Khulna [10] also found that about 80% of fecal sludge from on-site pit latrines is not safely managed [11].

Limited studies have been conducted to quantify levels of fecal contamination in different environmental compartments in urban Dhaka [12–15]. Direct ingestion of fecal contamination through contaminated drinking water has been studied extensively both at household and community levels in urban Bangladesh by measuring fecal indicator bacteria [16–19]. Other exposure pathways in urban Bangladesh, including contaminated soil [13], market produce [12], and street food [14] have been linked to adverse health outcomes such as diarrhea, environmental enteric dysfunction, and stunting [20,21]. Yet the contribution of these pathways to total fecal exposure remains understudied. Most urban studies have had small sample sizes, studied few communities, and targeted only a limited number of specific environmental compartments (i.e., market produce, soil, or street food), which are unlikely to provide a complete picture of the environmental fecal contamination levels in those communities. To inform evidence-based decision-making processes, policymakers, local government administrators, and local NGOs need data on the full range of fecal contamination pathways in order to more effectively prioritize and target interventions.

The SaniPath Exposure Assessment Tool quantitatively assesses exposureto fecal contamination via multiple pathways using a combination of microbiological data on environmental samples and information on the frequency of behaviors involving exposure to each environmental pathway [20,21]. We conducted a cross-sectional study to investigate the levels of environmental fecal contamination in different environmental compartments in 10neighborhoods in urban Dhaka using the SaniPath Exposure Assessment Tool [20]. In addition, information on relevant physical and demographic characteristics of the study neighborhoodswas collected.

## Methods

### Enrollment of study neighborhoods

Before study site selection, we conducted a stakeholder meeting and shared our protocol with local collaborators, partners, policymakers, and national and international NGOs to developneighborhood selection criteria based on the water and sanitation context in urban Dhaka. We selected neighborhoods based on socio-economic status, stability of the population (i.e., permanent vs. floating/transient population), nature of the housing and WASH infrastructure and services (i.e., unstructured vs. structured slums, and non-slums with poor WASH facilities/services, and non-slums with improved WASH facilities/services) and varied geographic locations [Dhaka South City Corporation (DSCC) and Dhaka North City Corporation (DNCC)] (Figure S1).

In 2011, the Dhaka City Corporation (DCC) was divided and re-created as DSCC and DNCC under an amendment act [22]. In this study, we collected an equal number of samples from neighborhoods in each corporation to explore differences in *E. coli* concentrations between the city corporations. We selected 10 neighborhoods from urban Dhaka (five from each city corporation) between April and June 2017: four low-income neighborhoods, two “floating” communities with transient populations, and four middle-to high-income neighborhoods (Table S1). We used the Bangladesh Bureau of Statistics (BBS) 2014 [23] slum list to select the low-income neighborhoods and floating communities for this assessment. We enrolled low-income neighborhoods that included at least 300 household compounds each and categorized them into “structured” (Kalshi and Shampur) and “unstructured” (Badda and Hazaribagh) slums. “Structured” slums had permanent household structures, >20 hours water supply per day, and shared latrine facilities. “Unstructured” slums had poorly structured housing (woods, bamboo, tin etc.), poor water distribution systems (i.e., through flexible pipes,) and poor sanitation facilities (i.e., mostly hanging toilets) compared to structured slums. We selected the Gabtoli bus terminal and Kamalapur railway station areas as floating communities because of the transient populations who live in these areasand do not have permanent dwellings. Four high- and middle-income communities were selected from two separate elite communities (Gulshan and Dhanmondi), one commercial/business area (Motijhil), and one newly developed neighborhood (Uttarkhan) from urban Dhaka (Figure S1 and Table S1).

In each neighborhood, we conducted one key informant interview (KII) with either a city official (i.e. city corporation staff and ward commissioners) or a community leader (i.e., local political leaders, religious leaders, or NGO workers/representatives) who had lived or worked in the selected neighborhood for more than five years and had a good understanding about the water, sanitation, and hygiene (WASH) facilities and practices of the neighborhood.

A total of 1,000 environmental samples (10 neighborhoods x 10 sample types x 10 samples per type) were collected. The sample types included: 1) swabs from the walls and door handles of shared/communal and/or public latrines accessed by any neighborhood residents, 2) soil/sand/mud frompublic areas where people gather and children commonly play, 3) open drain water from an open channel, carrying liquid and solid waste, including rainwater, floodwater, and wastewater from toilets and household activities, from locations where community people and children commonly come into contact, 4) bathing water from both municipal and non-municipal water supplies, 5) municipal drinking water (both legal and illegal connections) supplied by Dhaka Water Supply and Sewerage Authority (WASA) and accessed through piped water into compounds (including flexible pipes) and public taps/stand posts that are provided by the government or managed by someone in the community, 6) non-municipal drinking water (20 L commercially available jars or submersible pumps connected to a deep borehole), 7) surface water from community ponds and/or lakes, 8) floodwater that remains stagnant for at least one hour after rain, 9) produce that were commonly eaten raw, and 10) street food that was sold on the street and commonly consumed by community members including children (Table S2). We considered these to be priority environmental samples based on: 1) self-reported behavior about contact and ingestion from people in the study neighborhoods, 2) likelihood of contamination, as suggested by previous research in Bangladesh [12,14,15,17,18,24–27], 3) recommendations from the stakeholders meeting, and 4) information from the KIIs.

### Sampling siteselection

Before sample collection, the fieldworkers conducted a transect walk within each neighborhood and noted possible sampling sites for each type of sample in all the neighborhoods. In brief, for latrine swabs, fieldworkers purposively selected 10 shared/public latrines within each neighborhood that met the inclusion criteria. If there were multiple latrines in a latrine block, field workers selected the latrine thatwas reported most frequently used. Fieldworkers collected 10 soil samples in each neighborhood where children usually play. They also collected information on the type of soil (soil/sand/mixed), distance between the closest latrineand the sample site, and feces visible around the sampling area. For municipal and non-municipal drinking and bathing water, the fieldworkers first purposively selected 10 shared water points of each sample type in each neighborhood. Then, they recorded the source of the supplied water, type of connection (legal/illegal) and secondary extraction source (shallow tubewell, deep tubewell, public tap/standpipe, or piped water into the compound). Fieldworkers also measured the turbidity (LaMotte Model 2020i, LaMotte Company, Chestertown, MD) and/or free chlorine residual (LaMotte Model 1200, LaMotte Company, Chestertown, Maryland) of the water and recorded thevalues using a mobile device. For drain water samples, fieldworkers explored all open drains within the neighborhood during the transect walk and purposively selected 10 open drains where children play or people came in to contact with the drain water while walking. Floodwater samples were collected during the early monsoon (from June 1 to June 17, 2017). The fieldworkers collected the stagnant water that remained for at least one hour after rainingfrom the street and/or courtyard where children play or people came in to contact with the floodwater. Surface water samples were collected from the rivers, ponds, ditches, and/or lakes within the neighborhoods where children often swim or play or people wash utensils/clothing. During the transect walk, the fieldworkers explored all surface water sources in each neighborhood and purposively selected 10 sources geographically distant from each other. If the surface water source was small (pond/ditch), then the fieldworkers collected a single water sample, and if the water source was large (lake/river), the fieldworkers collected multiple samples from different points of the same source. We collected prepared street foods from street food vendors and/or from the street food shops depending on the availability during each day of sample collection. Food items sold on the streets and commonly eaten by the children and adults living in the community were collected. For this study, we collected *Fuska* (a round puffed and fried pastry with a hole on the top to fill with spiced sauce), c*hotpoti* (popular hot and sour snacks made of potatoes, chickpeas, onions, and chilies mixed with tamarind sauce), and *jhalmuri* (mixture of puffed rice and a variety of spices including peanuts, mustard oil, chili, onion, tomato, fresh ginger, salt, and lemon juice) (Table S2) [15,28]. For produce, the fieldworkers visited the local produce market in each neighborhood and sampled fresh produce that people commonly consumed raw or uncooked such as salad or garnish. Salads are typically prepared with bare hands and consist of raw vegetables like tomatoes, cucumbers, carrots, lettuce, coriander, onion, and green chili [29]. For this study, we collected samples of tomatoes, cucumbers, and coriander leaves, which are common salad ingredients found in Dhaka food markets.

### Environmental sample collection technique

We used SaniPath standard protocols [30] to collect all environmental samples except for street food which was not assessed during the previous SaniPath Tool assessments. After obtaining consent, the fieldworkers requested the street food vendors to prepare a single serving as he/sheusually prepares it. The fieldworker held a 500 mL Whirl-Pak bag (Nasco, FortAtkinson, WI) with the mouth open, and the vendor poured/placed the food into the bag.

After each sample was collected, fieldworkers sealed the bag, noted the time of sample collection, and immediately placed it into a cold box that was maintained at < 10°C with ice packs. Then, they used a mobile phone to record the Global Positioning System (GPS) coordinates of the sampling site and take at least two photographs of the sample and/or sampling site.

### Laboratory sample processing

A laboratory supervisor received the environmental samples within 4 hours of collection and analyzed the samples for *E. coli* using the IDEXX-Quanti-tray^®^ 2000 technique with Colilert-24 media (IDEXX Laboratories, Westbrook, Seattle, WA) [31] to quantify the most probable number (MPN) of *E. coli* per unit of sample. *E. coli* is commonly used as an indicator of fecal contamination in water, food, and environmental samples [13,32,33]. We chose to use *E. coli* to allow for comparison with other studies.

### Enumeration of *E. coli*

All environmental samples were processed on the same day, typically within 6 hours of collection, using the IDEXX Quanti-Tray 2000 system and Colilert reagent (IDEXX Laboratories, Maine, USA). Initially, different dilutions of samples were pre-tested to determine the ideal dilution factor to minimize samples with undetectable *E. coli* or *E. coli* exceeding the Quanti-Tray upper detection limit. Due to the wide range of the sampling sites (high income vs. low-income vs. floating),at least two dilutions per sample were analyzed to optimize detectionof positive *E. coli* wells within the Quanti-Tray detectable range of >1 to <=2419.6 MPN per tray (See supplemental information and Table S5 for detailed dilution procedures).

One field blank of distilled water was collected and processed each day. The laboratory technician filled one 100 mL Whirl-Pak bag with distilled water in the study community as a measure of the staff’s sterile technique. This blank was then tested in the laboratory for *E. coli*, and if the field blank showed any growth, we considered that contamination had occurred during sample collection and reinforced aseptic precautions for subsequent sample collection. Less than 1% of the tested blanks had positive growth. One laboratory blank per laboratory assistant per day, one positive control (drain water), and one negative control (distilled water) per batch of Colilert per laboratory assistant per day were processed for quality control. Finally, 100 mL environmental samples were processed and sealed in a Quanti-Tray and incubated at 37°C for 24 hours. The MPN of *E. coli* was determined by counting the number of fluorescing wells and calculating according to the manufacturer instructions. All water samples were reported as MPN of *E.coli*/100 mL, latrine swabs were reported as MPN of *E. coli*/swab, produce were reported as MPN of *E. coli*/single serving, and street food samples were reported as MPN of *E. coli*/gram.

### Qualitative data analysis

The fieldworker who recorded KIIs transcribed them in Bengali so that thematic content analysis could be performed [34]. The investigator manually coded the transcripts in an Excel spreadsheet according to the research objectives. After coding, the investigator categorized the data under different themes and matched these themes to factors associated with selection of environmental samples in each community.

### Quantitative data analysis

We substituted the value of 0.5 MPN for samples below the detection limit and 2419.6 MPN for samples above the detection limit, and calculated the *E. coli* concentration with corresponding dilution factors (Table S5). When the *E. coli* counts of all three dilutions were <1 MPN, we used the lowest diluted sample to estimate the concentration. When the *E. coli* counts of all three dilutions were >2419.6 MPN, we used the highest dilution to estimate the concentration, and if at least one *E. coli* count was within the detectable limit (from 1 to 2419.6 MPN) we calculated the average concentration of *E. coli* ignoring the censored (out of detectable limit) *E. coli* counts. *E. coli* concentrations were log_10_transformed, and summarized by sample type and neighborhood. We compared *E. coli* contamination between the low-income, high-income and floating neighborhoods (Table S1), and between the north and south parts of the city (DNCC and DSCC) using generalized linear regression models. We also examined differences in the level of contamination between neighborhoods graphically using an error bar graph produced by R (version 3.4.1). All statistical analyses were conducted using STATA-13.

## Results

### Key informant interviews (KII)

City officials, and/or community leaders reported that shared latrines were the most common type of latrine used by all communities. Key informants reported that open, rather than closed, drains were common in all neighborhoods except Dhanmondi. The municipal water supply was reported as the most common source of bathing and drinking water throughout the neighborhoods except for Kamalapur and Uttarkhan. In Kamalapur, most people used water from deep bore wells and commercially available 20L jar water, and in Uttarkhan, private submersible pumpsconnected to a deep borehole were the main source of drinking water. Commercially available jar waterwas also reported as the most commonly used drinking water in all neighborhoods except for Uttarkhan. Almost all city officials, and/or community leaders reported that *fuska, chotpoti* and *jhalmuri* were commonly eaten street foods and that cucumbers, tomatoes, and coriander were commonly eaten raw vegetables in all neighborhoods (Table S3).

### Magnitude of *E. coli* contamination in environmental samples

Among environmental samples, almost all drain water (98%) and street food (93%) samples, nearly 80% of fresh produce, surface water and floodwater samples, and more than 50% of municipal drinking water, non-municipal drinking water and bathing water samples were contaminated with *E. coli* (Figure 1). Characteristics of individual samples are described in supplemental Table S4.

**Figure 1:**
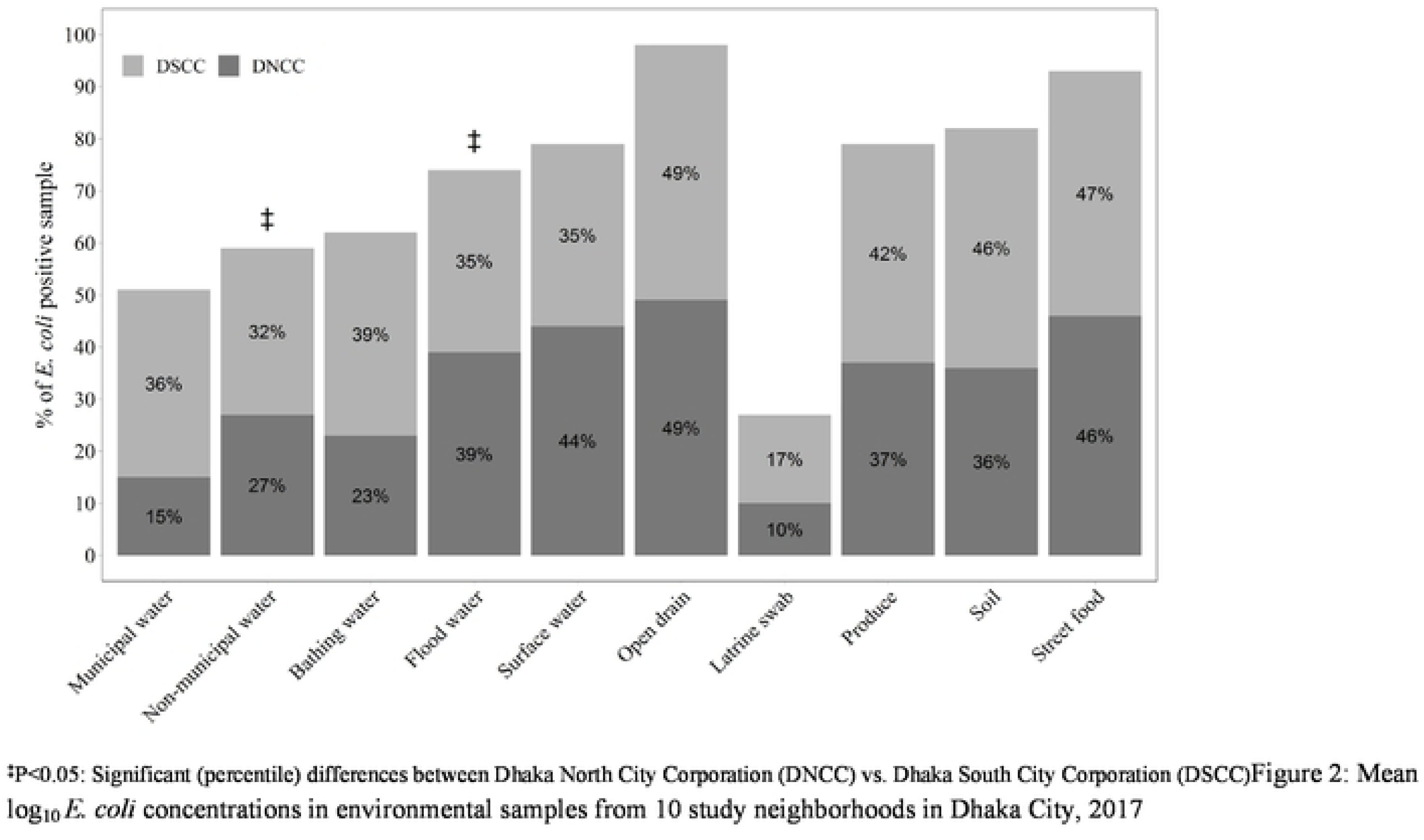
Percentage* of *E. coli* positive environmental samples [(N= 704, DSCC =378, DNCC=326)] from 10 study neighborhoods in Dhaka city, 2017

**Figure 2:**
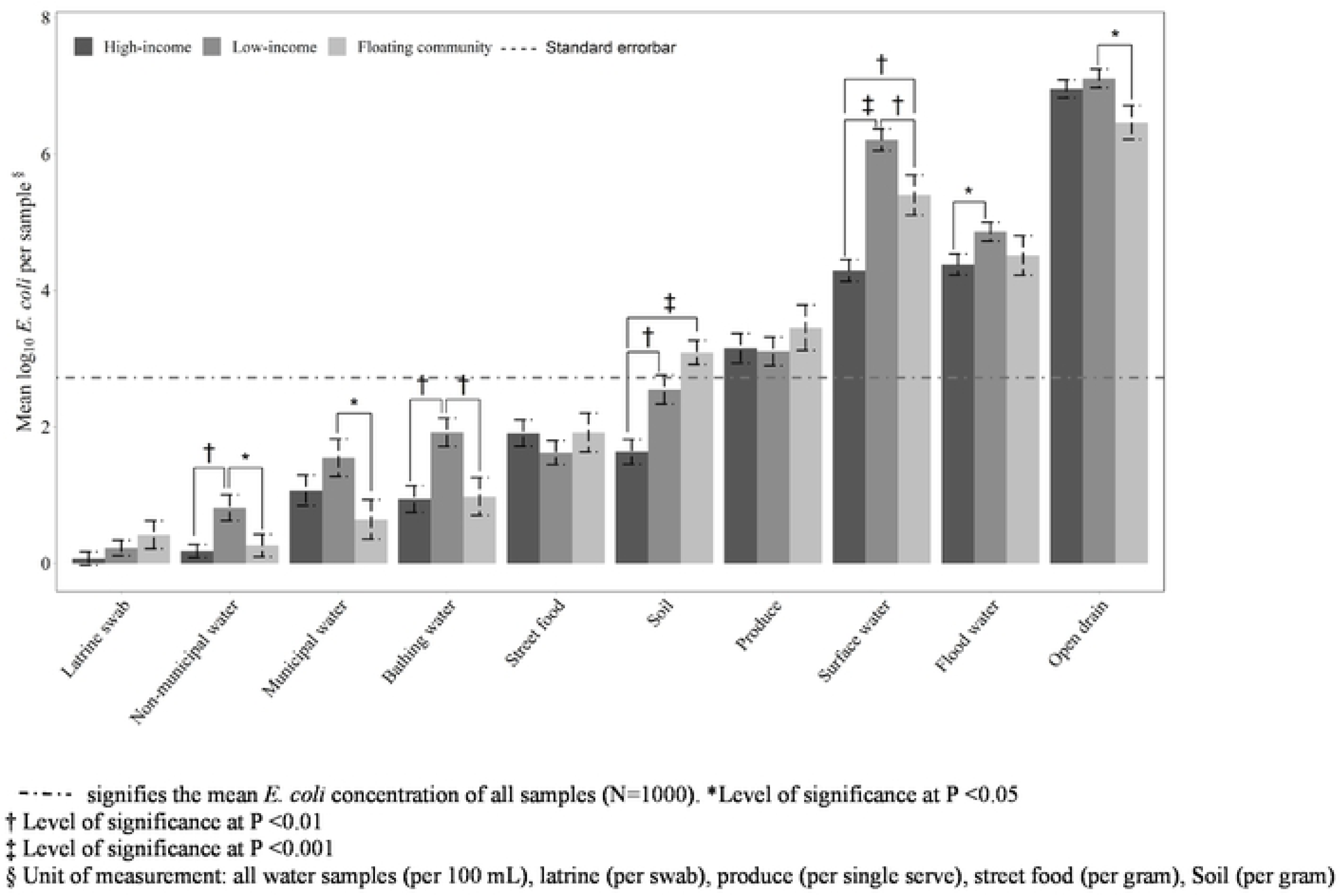
Mean log_10_ *E. coli* concentrations in environmental samples from 10 study neighborhoods in Dhaka City, 2017

Among the 10 neighborhoods, Hazaribagh had the greatest concentration of *E. coli* [mean (SD)] in five categories of samples, including shared latrine swabs [0.64 log_10_ MPN/swab (0.99)], municipal drinking water [3.20log_10_ MPN/100 mL (0.84)], non-municipal drinking water [1.67 log_10_ MPN/100 mL (1.21)], surface water [7.38 log_10_ MPN/100 mL (0.00] and floodwater [5.47 log_10_ MPN/100 mL (0.89)]). *E. coli* concentrations in drain water [7.61log_10_ MPN/100 mL (0.65)] from Shampur, street food [2.58log_10_ MPN/gram (1.36)] from Dhanmondi, soil [3.19 log_10_ MPN/gram (1.49)] from Badda, produce [3.52 log_10_ MPN/serving (1.45)] from Kamalapur, and bathing water [2.57 log_10_ MPN/100 mL (1.57)] from Shampur were higherthan other neighborhoods.The concentration of *E. coli* in shared latrine swabs, bathing and municipal water and street food from Gulshan, and soil, non-municipal drinking water, surface water and floodwater samples from Uttarkhan were the lowest compared to rest of the neighborhoods (Table 1).

**Table 1:** Mean log_10_ *E. coli* concentration in environmental samples from 10 Dhaka city neighborhoods, 2017

| Neighborhoods | Environmental Samples, Mean log<sub>10</sub>MPN *E. coli* concentration (SD) |  |  |  |  |  |  |  |  |  |
| --- | --- | --- | --- | --- | --- | --- | --- | --- | --- | --- |
|  | Latrine swab<sup>b</sup> | Soil<sup>d</sup> | Drain water<sup>a</sup> | Bathing water<sup>a</sup> | Municipal drinking water<sup>a</sup> | Non-municipal water<sup>a</sup> | Surface water<sup>a</sup> | Produce<sup>c</sup> | Street food<sup>d</sup> | Flood water<sup>a</sup> |
| N=10 Samples/neighborhood |  |  |  |  |  |  |  |  |  |  |
| **Floating communities** |  |  |  |  |  |  |  |  |  |  |
| Gabtolli (N) | 0.04<br>(0.63) | 3.11<br>(1.02) | 6.69<br>(0.97) | 0.33<br>(0.87) | 0.12<br>(0.64) | -0.15<br>(0.21) | 5.77<br>(1.32) | 3.37<br>(1.60) | 1.79<br>(1.55) | 4.21<br>(1.64) |
| Kamalapur (S) | 0.79<br>(1.02) | 3.06<br>(0.53) | 6.23<br>(1.23) | 1.62<sup>‡</sup><br>(1.24) | 1.16<br>(1.59) | 0.67<sup>§</sup><br>(0.84) | 5.02<br>(1.28) | **3.52**<br>(1.45) | 2.04<br>(1.0) | 4.81<br>(0.74) |
| **Unstructured slums** |  |  |  |  |  |  |  |  |  |  |
| Kalshi (N) | 0.05<br>(0.64) | 2.26<br>(1.49) | 6.64<br>(0.78) | 2.12<br>(0.89) | 0.16<br>(0.57) | 0.69<br>(1.27) | 6.05<br>(0.74) | 2.57<br>(1.23) | 1.44<br>(1.27) | 4.57<br>(0.57) |
| Shampur (S) | -0.06<br>(0.29) | 2.00<br>(1.09) | **7.61**<sup>§</sup><br>(0.65) | **2.57**<br>(1.57) | 2.71<br>(1.55) | -0.19<sup>‡</sup><br>(0.34) | 5.86<br>(0.92) | 3.55<br>(1.59) | 0.76<br>(0.86) | 4.52<br>(1.15) |
| **Structured slums** |  |  |  |  |  |  |  |  |  |  |
| Badda (N) | 0.26<br>(0.65) | **3.19**<br>(1.49) | 6.76<br>(1.10) | 0.71<br>(0.92) | 0.11<br>(0.72) | 1.09<br>(0.93) | 5.55<br>(0.92) | 3.46<br>(1.25) | 2.26<br>(1.02) | 4.88<br>(0.39) |
| Hazaribag (S) | **0.64**<br>(0.99) | 2.82<br>(1.01) | 7.43<br>(0.41) | 2.28<sup>‡</sup><br>(1.06) | **3.20**<br>(0.84) | **1.67**<br>(1.21) | **7.38**<br>(0.01) | 2.85<br>(1.06) | 2.02<br>(0.66) | **5.47**<br>(0.89) |
| **Non-slum with poor WASH** |  |  |  |  |  |  |  |  |  |  |
| Uttarkhan (N) | 0.36<br>(0.96) | 1.08<br>(1.36) | 7.00<br>(0.62) | 1.07<br>(1.21) | 1.96<br>(1.62) | -0.12<br>(0.25) | 4.51<br>(1.29) | 2.87<br>(1.29) | 1.79<br>(1.21) | 4.02<br>(0.68) |
| Motijheel (S) | 0.12<br>(0.76) | 2.29<sup>‡</sup><br>(0.60) | 6.68<br>(1.00) | 1.62<br>(1.10) | 1.57<br>(1.02) | 0.32<br>(0.69) | 3.43<sup>‡</sup><br>(0.71) | 3.14<sup>‡</sup><br>(1.18) | 1.92<br>(1.05) | 4.81<sup>‡</sup><br>(0.75) |
| **Non-slum with improved WASH** |  |  |  |  |  |  |  |  |  |  |
| Gulshan (N) | -0.15<br>(0.01) | 1.57<br>(1.30) | 7.01<br>(0.49) | 0.18<br>(0.87) | -0.47<br>(0.71) | 0.01<br>(0.48) | 5.07<br>(0.64) | 3.23<br>(1.47) | 1.34<br>(1.05) | 4.62<br>(0.38) |
| Dhanmondi (S) | -0.05<br>(0.26) | 1.60<br>(1.02) | 7.15<br>(1.14) | 0.89<br>(1.45) | 0.82<br>(1.33) | 0.50<br>(0.72) | 4.16<sup>§</sup><br>(0.53) | 3.36<br>(1.66) | **2.58**<sup>‡</sup><br>(1.36) | 4.07<br>(1.56) |
| Mean log<sub>10</sub> *E. coli* in 10 study neighborhoods | 0.20<br>(0.73) | 2.29<br>(1.29) | 6.91<br>(0.92) | 1.34<br>(1.34) | 1.17<br>(1.55) | 0.45<br>(0.94) | 5.28<br>(1.37) | 3.19<br>(1.36) | 1.79<br>(1.18) | 4.60<br>(1.02) |
| Mean difference DNCC minus DSCC (95% CI) | -0.17<br>(-0.47, 0.11) | 0.09<br>(-0.60, 0.42) | -0.20<br>(-0.57, 0.16) | -0.91<br>(-1.42, -0.41) | -1.43<br>(-1.98, -0.89) | -0.29<br>(-0.66, 0.08) | 0.22<br>(-0.32, 0.77) | -0.18<br>(-0.72, 0.36) | -0.14<br>(-0.61, 0.33) | -0.27<br>(-0.68, 0.13) |

N=Dhaka North City Corporation (DNCC), S=Dhaka South City corporation (DSCC), <sup>‡</sup>Level of significance between neighborhoods at P <0.05, <sup>§</sup> Level of significance at P <0.01, <sup>||</sup> Level of significance at P <0.001, **Bold digits**: Highest mean log<sub>10</sub> *E. coli* concentration in specific type of environmental sample
<sup>a</sup>All water samples including drains were reported as MPN of *E. coli*/100 mL; <sup>b</sup>Latrine swabs were reported as MPN of *E. coli*/swab; <sup>c</sup>Produce were reported as MPN of *E. coli*/single serving; <sup>d</sup>Street food and soil samples were reported as MPN of *E. coli*/gram.

Although overall concentration of *E. coli* in most of the sample types were similar between DNCC and DSCC, *E. coli* concentrations were significantly higher in bathing water [log_10_ mean difference DNCC minus DSCC = −0.91 log_10_ MPN/100 mL (95% CI: −1.42, −0.41] and municipal drinking water [log_10_ mean difference DNCC minus DSCC = −1.43 log_10_ MPN/100mL (95% CI: −1.98, −0.89)] from DSCC compared to DNCC (Table 1 and Figure 1).

Overall, the municipal drinking water was more contaminated compared to non-municipal water (mean difference: non-municipal minus municipal water = −0.73 log_10_ MPN/100 mL, 95% CI: −1.08, −0.37). Although, the *E. coli* concentrations were similar between the municipal and non-municipal water in DNCC, the *E. coli* concentration was significantly higher in municipal water in DSCC (mean difference: non-municipal drinking water minus municipal drinking water = −1.30 log_10_ MPN/100 mL, 95% CI: −1.81, −0.79) (Table 2). As expected, samples of municipal water generally had lower concentrations of *E. coli* than bathing water, floodwater, surface water and drain water.

**Table 2:** Differences between *E. coli* concentrations per 100 mL in samples from municipal drinking water and other types of water in 10 neighborhoods in Dhaka city, 2017.

| Neighborhoods | Mean log<sub>10</sub> MPN *E. coli* concentration differences from different water samples vs. municipal drinking water (95% CI) |  |  |
| --- | --- | --- | --- |
|  | DSCC (N=50) | DNCC (N=50) | All 10 study neighborhoods<br>N=100) |
| Municipal drinking water<sup>‡§</sup> | 1.89<sup>‡</sup> | 0.46<sup>‡</sup> | 1.18<sup>‡</sup> |
| Drain water vs. municipal water | **5.12**<br>**(4.61, 5.64)** | **6.36**<br>**(5.96, 6.75)** | **5.74**<br>**(5.39, 6.09)** |
| Bathing water vs. municipal water | -0.09<br>(-0.67, 0.48) | 0.42<br>(-0.33, 0.88) | 0.16<br>(-0.23, 0.57) |
| Floodwater vs. municipal water | **2.84**<br>**(2.31, 3.37)** | **4.00**<br>**(3.59, 4.10)** | **3.42**<br>**(3.06, 3.79)** |
| Non-municipal vs. municipal water | **-1.30**<br>**(-1.81, -0.79)** | -0.15<br>(-0.56, 0.25) | **-0.73**<br>**(-1.08, -0.37)** |
| Surface water vs. municipal water | **3.28**<br>**(2.66, 3.89)** | **4.93**<br>**(4.48, 5.39)** | **4.10**<br>**(3.69, 4.51)** |

\* Dhaka South City Corporation (DSCC), <sup>†</sup>Dhaka North City Corporation (DNCC)
<sup>‡</sup>Reference value: *E. coli* contamination in municipal drinking water
<sup>§</sup>The first row shows mean log<sub>10</sub> *E. coli* concentration and the other columns show the mean log<sub>10</sub> difference compared to municipal drinking water
<sup>||</sup>mean differences were significantly different between the two comparison groups

### Comparison of *E. coli* concentration across high-income, low-income, and floating neighborhoods

#### Low-income vs. high-income neighborhoods

There was no significant difference in *E. coli* contamination for latrine swabs, drain, municipal drinking water, produce and street foods between low-income and high-income neighborhoods (Table 3). The remaining five sample types had significantly higher *E. coli* concentrations in low-income neighborhoods [soil (mean difference:low-income minus high-income = 0.91 log_10_ MPN/gram, 95% CI: 0.39, 1.43), bathing water (mean difference:low-income minus high-income = 0.98 log_10_ MPN/100 mL, 95% CI:0.41, 1.54), non-municipal water (mean difference:low-income minus high-income = 0.64 log_10_ MPN/100 mL, 95% CI, 0.24, 1.04), surface water (mean differences from low-income minus high-income = 1.92 log_10_ MPN/100 mL, 95% CI: 1.44, 2.40) and floodwater (mean difference: low-income minus high-income = 0.48 log_10_ MPN/100 mL, 95% CI: 0.03, 0.92)] (Table 3).

**Table 3:** Comparisons between mean log_10_ MPN *E. coli* concentrations in environmental samples from low-income, high-income, and floating neighborhoods in Dhaka city, 2017.

| Neighborhoods | Latrine surface swab<sup>b</sup> | Soil<sup>d</sup> | Drain water<sup>a</sup> | Bathing water<sup>a</sup> | Municipal drinking water<sup>a</sup> | Non-municipal water<sup>a</sup> | Surface water<sup>a</sup> | Produce<sup>c</sup> | Street food<sup>d</sup> | Floodwater<sup>a</sup> |
| --- | --- | --- | --- | --- | --- | --- | --- | --- | --- | --- |
| Mean log<sub>10</sub> MPN *E. coli* concentration (SD) |  |  |  |  |  |  |  |  |  |  |
| Low-income* (N=40) | 0.22 (0.72) | 2.55 (1.34) | 7.11 (0.86) | 1.92 (1.32) | 1.55 (1.73) | 0.81 (1.19) | 6.21 (1.01) | 3.10 (1.13) | 1.62 (1.11) | 4.86 (0.88) |
| High-income† (N=40) | 0.07 (0.63) | 1.63 (1.16) | 6.95 (0.84) | 0.94 (1.23) | 1.07 (1.41) | 0.18 (0.60) | 4.29 (1.01) | 3.15 (1.37) | 1.91 (1.21) | 4.38 (0.98) |
| Floating‡ (N=20) | 0.42 (0.91) | 3.09 (0.79) | 6.46 (1.11) | 0.99 (1.24) | 0.64 (1.30) | 0.26 (0.73) | 5.40 (1.32) | 3.45 (1.49) | 1.92 (1.28) | 4.51 (1.28) |
| Mean log<sub>10</sub> MPN *E. coli* concentration differences between neighborhoods (95% CI) |  |  |  |  |  |  |  |  |  |  |
| Low-income vs. high-income | 0.15<br>(-0.17, 0.48) | **0.91**<br>**(0.39, 1.43)**<sup>§</sup> | 0.16<br>(-0.25, 0.56) | 0.98<br>(0.41, 1.54)<sup>§</sup> | 0.48<br>(-0.20, 1.16) | **0.64**<br>**(0.24, 1.04)**<sup>§</sup> | **1.92**<br>**(1.44, 2.40)**<sup>§</sup> | -0.04<br>(-0.65, 0.56) | -0.29<br>(-0.81, 0.24) | **0.48**<br>**(0.03, 0.92)**<sup>§</sup> |
| Low-income vs. floating | -0.19<br>(-0.59, 0.20) | -0.54<br>(-1.18, 0.95) | **0.65**<br>**(0.15, 1.14)**<sup>§</sup> | **0.94**<br>**(0.25, 1.63)**<sup>§</sup> | **0.91**<br>**(0.07, 1.73)**<sup>§</sup> | **0.55**<br>**(0.06, 1.04)**<sup>§</sup> | **0.81**<br>**(0.22, 0.40)**<sup>§</sup> | -0.35<br>(-1.09, 0.40) | -0.29<br>(-0.94, 0.34) | 0.35<br>(-0.20, 0.89) |
| Floating vs. high-income | 0.35<br>(-0.05, 0.75) | **1.46**<br>**(0.82, 2.10)**<sup>§</sup> | -0.49<br>(-0.98, -0.00) | 0.03<br>(-0.65, 0.72) | -0.43<br>(-1.26, 0.40) | 0.08<br>(-0.41, 0.57) | **1.10**<br>**(0.52, 1.69)**<sup>§</sup> | 0.30<br>(-0.44, 1.04) | 0.01<br>(-0.64, 0.65) | 0.13<br>(-0.42, 0.67) |

\*Low-income neighborhoods: Kalshi, Shampur, Badda, and Hazaribagh; †High-income neighborhoods: Gulshan, Dhanmondi, Motijhil, and Uttarkhan; ‡Floating neighborhoods: Gabtoli and Kamalapur neighborhoods; § mean differences were significantly different between the two comparison groups
<sup>a</sup>All water samples including drains were reported as MPN of *E. coli*/100 mL; <sup>b</sup>Latrine swabs were reported as MPN of *E. coli*/swab; <sup>c</sup>Produce were reported as MPN of *E. coli*/single serving; <sup>d</sup>Street food samples were reported as MPN of *E. coli*/gram.

#### Low-income vs. floating neighborhoods

Although concentrations of *E. coli* between low-income and floating neighborhoods were similar for latrine swabs, soil, produce, street food and floodwater samples, the concentrations were significantly higher in samples from low-income neighborhoods compared to floating neighborhoods for drain water(mean difference: low-income minus floating neighborhoods = 0.65 log_10_ MPN/100 mL, 95% CI: 0.15, 1.14), bathing water (mean difference: low-income minus floating neighborhoods = 0.94 log_10_ MPN/100 mL, 95% CI: 0.25, 1.63), municipal water (mean difference: low-income minus floating neighborhoods = 0.91 log_10_ MPN/100 mL, 95% CI: 0.07, 1.73), non-municipal water (mean difference: low-income minus floating neighborhoods = 0.55 log_10_ MPN/100 mL, 95% CI: 0.06, 1.04) and surface water (mean difference: low-income minus floating neighborhoods = 0.81 log_10_ MPN/100 mL, 95% CI: 0.22, 1.40) (Table 3).

#### Floating vs. high-income neighborhoods

We found similar *E. coli* concentrations between floating and high-income neighborhoods for all environmental sample types except for soil (mean difference floating minus high-income neighborhoods: 1.46 log_10_ MPN/gram, 95% CI: 0.82, 2.10) and surface water (mean differencefloating minus high-incomeneighborhoods = 1.10 log_10_ MPN/100 mL, 95% CI: 0.52, 1.69) (Table 3).

## Discussion

Extensive *E. coli* contamination was detected in most of the environmental samples collected throughout the 10 urban study neighborhoods, suggesting that all residential areas of Dhaka may be prone to fecal contamination regardless of geographic location or socio-economic status. This is consistent with the prediction of the fecal waste flows analysis for Dhaka [11,35] that estimated that 98-99% of fecal waste in Dhaka is ultimately distributed within the urban environment – including residential areas. Few studies have attempted to comprehensively measure fecal contamination in urban Dhaka. Previous studies have focused only on specific pathways, but they have also reported high occurrence of fecal contamination in environmental samples. A recent study in a large wholesale produce market and neighborhood retail markets in Dhaka found that 100% of carrot and red amaranth rinses, 92% of eggplant rinses, and 46% of tomato rinses were contaminated with *E. coli* [12]. Street-vended foods in Dhaka markets [14] and near schools (60% *jhalmuri*, 29% *chotpoti*) [15] were also reported to be highly contaminated with fecal bacteria. The detection of fecal indicator bacteria in these environmental samples suggests the potential presence of pathogenic organisms and the potential risk of enteric disease among Dhaka residents who are frequently exposed to these contaminated environments, drink contaminated municipal water, and/or consume raw or undercooked produce or street foods [36].

Unlike previous SaniPath deployments [20] that focused primarily on low-income neighborhoods, the Dhaka SaniPath assessment compared environmental contamination in a range of high-income, low-income, and floating communities. This diversity of neighborhoods allowed examination of fecal contamination that may be due to localized sources, such as a contaminated surface water body, vs. fecal contamination that moves through the city among both poor and wealthy neighborhoods through vehicles such as contaminated produce, municipal piped water, or open drains. Our results suggest that, despite socio-economic and infrastructure differences between the study neighborhoods, the fecal contamination levels for some sample types, like drain water, municipal drinking water, produce, and street food,were similar across neighborhoods. The widespread fecal contamination in these urban neighborhoods may be due to unsafe fecal sludge management and consequent movement and distribution of fecal contamination in the urban environment (i.e., through flooding, poor drainage systems, and/or unsafe dumping of sludge) [3]. Previous analyses of existing sanitation data concluded that <1% of household fecal sludge in Dhaka was effectively managed, and the vast majority of waste water and fecal sludge was not contained and was either leaking out of pipes and latrines or deliberately discharged directly into the environment [35]. Our primary data collection confirms the presence of fecal contamination in the range of residential environments that we studied.

Conversely, non-municipal drinking water, bathing water, surface water, and soil samples had significantly higher *E. coli* concentrations in low-income neighborhoods compared to high-income neighborhoods and suggests that the contamination in these pathways may be due to local sources of fecal discharge. Low-income urban neighborhoods are located mainly in lower elevations and in the periphery of the city (i.e., Hazaribagh) [37], where flooding occurs almost every year [38]. The floodwater runs off into storm sewers and ultimately into surface water, and during heavy rainfall, the contaminated water returns to the environment and contaminates the soil [39]. Poor drainage systems, improper child feces disposal, and poor fecal sludge management likely increase the fecal contamination of the soil in low-income neighborhoods [40]. Lastly, unimproved housing infrastructure (i.e., dirt floor/walkway), poor hydraulic and physical integrity of the water distribution network (leaky flexible pipes and illegal connections), unsafe water storage, high population density, and poorly designed and constructed on-site household and community sanitation systems that do not adequately contain fecal sludge may contribute to higherlocalized fecal contamination levels in soil and water in low-income neighborhoods in Dhaka [41,42].

The overall municipal water quality results reported here are consistent with previous studies in Dhaka that reported high levels of fecal contamination in municipal drinking water mostly in low-income communities [17,43]. A nationally-representative water quality assessment estimated that 41% of all improved water sources sampled across Bangladesh were contaminated with *E. coli* [44,45]. Piped water systems, which are almost exclusive to urban areas of Bangladesh, were among the most contaminated drinking water sources. That assessment also reported that 55% of the water samples from municipal public taps and more than 80% of the samples from water taps on premises in urban neighborhoods of Bangladesh had *E. coli* contamination [11]. Contamination can occur either in the distribution system due to frequent pipe breaks and illegal connections, low or negative water pressure due to intermittent service, and/or because of poor domestic water storage structures and maintenance [25,46,47].

Although fecal contamination was widespread throughout urban Dhaka, we found significantly higher concentrations of *E. coli* in most of the samples from DSCC compared to DNCC. There are several possible explanations for these differences. Firstly, DSCC is an older part of the city with older infrastructure (i.e., pipes, drainage) and narrow lanes that largely lack a drainage system. These lanes often become flooded during rainfall [70]. Additionally, the households of DSCC are closely packed together with a leaky water distribution system and older sanitation facilities. Furthermore, the population density is about three times greater in DSCC (>124,000 persons per square kilometer) compared to DNCC (<35,000 persons per square kilometer) [71], and this presents an additional challenge to ensure adequate WASH services in DSCC with limited resources. Finally, the highly polluted Buriganga River passes beside DSCC and is a major source of environmental contamination.

Our results show that the low-income communities in DSCC had significantly higher *E. coli* concentrations in their municipal water supply compared to the low-income communities of DNCC (Table 1). In DSCC, the majority of the municipal water is distributed through the Saidabad surface water treatment plant and from deep bore wells, and in DNCC, water is exclusively supplied by deep bore wells [49]. The high concentration of *E. coli* in DSCC municipal water supply may be due to the long water residence time in a water distribution system with compromised physical and hydraulic integrity that allows intrusion of contamination. Additionally, recent research on water quality in low-income urban communities in Dhaka reported that most of the municipal water sources do not have chlorine injectors and/or that the water was inappropriately treated before distribution [17,43].

High concentrations of fecal contamination have frequently been reported on produce in low- and middle-income countries, including Bangladesh [12] and elsewhere [20,24,50–53]. Fresh produce can be a vehicle for fecal contamination to move across the city to both poor neighborhoods and high-income households [20] and can pose a major health risk to urban populations [54]. Limited data are available on disease burdens attributed to food contamination in low- and middle-income countries [55,56]. The CDC estimates that nearly half of all food-borne illnesses in the United States [57] are caused by contaminated fresh produce and that more than 30% of gastroenteritis cases in low- and middle-income countries are linked to food borne transmission [58]. Risk factors for food contamination in the low- and middle-income settings are different than high-income countries because cooked food is more common and is usually freshly prepared in households [29,59]. The causes of the fecal contamination detected on the produce in this study are not known and may be due to poor agricultural practices by farmers (e.g. use of wastewater for irrigation) and unhygienic conditions in the produce markets. Observational studies in rural Bangladesh identified that produce washing practices during salad preparation (uncooked and mashed cucumber, tomato etc.) within low-income neighborhoods were inadequate, and salads were often contaminated due to poor hygiene practices [29,60].

Over 90% of street food samples in this study were contaminated with *E. coli*, and there was no geographical variation in the level of contamination. This is a major public health concern, and a number of studies have reported that people who patronize street food vendors suffer from food-borne diseases like diarrhea, cholera, typhoid fever, and other enteric diseases [36,61,62]. A number of studies in Bangladesh [6,15,61,63,64] and elsewhere [36,62,65,66] also found high levels of microbial contamination in street-vended foods. These foods can be contaminated in different ways. According to a government report, 94% of street food vendors in Dhaka reported that they used the municipal water supply to prepare food and did not take any measures to treat the water. The report also found that nearly 58% of the vendors did not cover their food while selling and most vendors did not wash their hands with soap while preparing the food [6]. Additionally, most of the vendors (68%) were located on footpaths; 30% of vending carts were placed near drains; and 18% were placed near sewerage.

### Strengths and Limitations

This study is the most comprehensive and systematic assessment of fecal contamination in urban Dhaka ever conducted and included not only a wide range of neighborhoods but also examined 10 different types of environmental samples for fecal indicator bacteria. While this study provides valuable information on both the magnitude of fecal contamination in the environment and how it is distributed in the city, it also has some important limitations. First, although a large number of environmental samples were collected from three types of neighborhoods with different socio-economic status in an attempt to represent a range of conditions, it was not possible to cover the entire city. Therefore, our findings may not be generalizable to all urban neighborhoods in Dhaka, in Bangladesh, or to other cities in South Asia, such as those with dry climates or with better fecal sludge management and improved WASH facilities [20]. Additionally, while the sample size was appropriate for the primary study objective of conducting an exposure assessment, it may not be sufficient for detecting modest differences between individual neighborhoods or between environmental pathways. Future studies of environmental contamination should increase the sample size for pathways that have large variation and/or cover larger or more diverse geographical regions [72].

In this study, we measured *E. coli* as fecal indicator bacteria but did not attempt to detect specific enteric pathogens in the environment – some of which survive longer than *E. coli* and are highly infectious even at low concentrations. We are not able to estimate the disease burden associated with the levels of fecal contamination that were detected in these neighborhoods. A recent study in Dhaka suggested that multi-drug resistant (MDR) *E. coli* were widespread in the public water supply in Dhaka, which could be potentially hazardous for human health [73]. Further, we did not distinguish if the source of *E. coli* we detected was from animals or humans. Although animal density in Dhaka neighborhoods is low, a recent study reported that a ruminant-associated bacterial target was detected in 18% of hand rinse and 27% of floor samples in a study neighborhood in Dhaka [13]. A review in 2017 also suggested that exposure to animal feces in urban environments may be associated with enteric diseases, soil-transmitted helminths infections, environmental enteric dysfunction, and growth faltering [74]. These findings suggest that effective community fecal management should account not only for human sources of contamination but also for animal sources in urban environments.

## Conclusions and Recommendations

The results of this study indicate that there is widespread fecal contamination in the public domain in Dhaka in both low-income and high-income neighborhoods. The poor drainage system, poor sanitation facilities, frequent flooding and poorly managed municipal water supply of Dhaka may contribute to this extensive fecal contamination [75].The evidence from this study can inform policies and interventions to protect public health in Dhaka and can also identify important research needs. Intervention strategies should consider how the geographic, infrastructure, and economic differences across the city impact various fecal exposure pathways and their implications for effectively reducing fecal contamination in urban neighborhoods of Dhaka.

The high prevalence of municipal drinking water contamination reported here emphasizes the importance of adopting appropriate organizational arrangements for the routine maintenance and improvement of drinking water systems in order to prevent contamination in the municipal piped network and alertingwater utilityand municipal authorities to problems with the system that need to be addressed [48]. Appropriate, affordable, and effective centralized, community-level, and household-level water treatment and storage technologies need to be developed along with increased awareness among landlords and compound managers about the importance of safe water management practices in both the public and private domain [48]. Future studies should examine the excessive water contamination detected in DSCC andidentify the specific factors that contribute to this problem.

Of special concern is the evidence that the food supply in the city (fresh produce and street-vended food) has high contamination levels and poses a risk citywide. This risk may be less visible than poor WASH infrastructure and therefore less targeted for intervention. Policies and regulations for safe street food are weak and poorly enforced in most low- and middle-income countries [67] and even non-existent in some countries [68] including Bangladesh [6]. Therefore, formulation of appropriate food hygiene policies and proper enforcement are essential to reduce therisks associated with street food consumption [65,69]. Further studies of the causes of food contamination at farms, markets, and street vendors are needed to understand the critical points in the food production chain where contamination is introduced and how to prevent this contamination and mitigate risk through changes in agricultural practices and food handling and hygiene.

Improving fecal sludge management, training on food hygiene and produce handling for food vendors, and improving the microbial quality of municipal water should be explored as strategies to prevent the introduction of fecal contamination into different environmental pathways. Long-term integrated programs that include the provision of urban WASH services, housing/infrastructure improvement, behavior change communication, appropriate technology development (i.e., safely-managed sanitation systems, online automated centralized and community-level water treatment systems), improved food safety practices, and good personal hygiene) [76], could reduce fecal contamination and improve the overall WASH conditions in urban neighborhoods. Future studies should explore behavior that brings people into contact with the environment and assess exposure to environmental fecal contamination through multiple pathways and the associated risks in different neighborhoods of urban Dhaka. Application of sound risk analyses and formulation of appropriate environmental protection policies are necessary to provide a strong scientific basis for the host of risk management options that Dhaka city authorities may need to explore in order to ensure public health and safety and achieve Sustainable Development Goal 6 (safely managed water and sanitation) by 2030.

## Acknowledgments

The study was financially supported by the Bill & Melinda Gates Foundation (grant no. 00010161) through the Rollins School of Public Health at Emory University. icddr,b acknowledges with gratitude the commitment of the Bill & Melinda Gates Foundation and Rollins School of Public Health, Emory Universityto its research efforts. We acknowledge theWorld Bank Bangladesh Country Office team for their efforts in partnering, workshops, and contributing to sampling/study design. We are grateful to James Michiel, Senior mHealth and Informatics Analyst for his support at the beginning of the project. We also acknowledge the efforts of the Dhaka SaniPath field team: The study design, data collection and data entry were conducted by the SaniPath project team: Badal Howlader, Khan Ali Afser, Rana Mia, Shamim Ahamed, Abdul Barek, Mohammad Rafik, Emdadul Haque, Raju Ahmed, and Md. Arifin. icddr,b is also grateful to the Governments of Bangladesh, Canada, Sweden and the UK for providing core/unrestricted support.

---

**Table S1:** Definitions of neighborhoods for SaniPath Dhaka deployment, 2017

| Definition of neighborhoods |  |
| --- | --- |
| Floating community (BBS 2014) [23] | Floating communities are made up of transient people who have no permanent dwelling units, and/or travelers from different parts of the country. These are the areas with the biggest bus terminals, railway stations and truck stands where thousands of people come every day and stay temporarily. (We selected Gabtoli bus stand and Kamalapur railway station areas as floating communities.) |
| Unstructured slum | A slum where most of the houses are constructed of very poor materials, such as walls and roofs made of straw, leaves, gunny sacks, polythene paper, bamboo, or tin. Most of the latrines are either hanging or made with bamboo, straw leaves, gunny sacks, and/or polythene paper. The majority of the water is supplied through flexible leaky pipes outside the compound or in the public space. (We selected Kalshi and Shampur areas as unstructured slum communities.) |
| Structured slum | A slum with moderately good household structures constructed from bricks and/or tin. Most of the households have access to shared piped drinking water within the compound and have relatively improved/shared latrine facilities compared to an unstructured slum. (We selected Badda and Hazaribagh areas as structured slum communities.) |
| Slum/low-income community (BBS 2014) [23] | A slum/low-income community is a cluster of compact settlements of five or more households that generally grows haphazardly and may be located on government or private vacant land. (For analysis, we combined both unstructured (Kalshi+Shampur) and structured slums (Badda+Hazaribagh) together for the category of slum/ low-income communities.) |
| High-income elite community | These are the affluent neighborhoods in Dhaka: a residential area for elites and home to a number of the city's restaurants, shopping centres, schools, and members' clubs. These neighborhoods also host the majority of embassies and high commissions in Dhaka. These areas include Gulshan, Baridhara, Banasree, Dhanmondi R/A, Bashundhara R/A, etc., and they attract upper-class residents. These areas also have improved WASH facilities compared to rest of the city. (We selected Gulshan and Dhanmondi areas as high-income elite communities) |
| Middle-income commercial/business area | This is a major business and commercial hub of Dhaka city and has more offices and business institutions than any other part of the city. It is home to the largest number of corporate headquarters in the nation. Government officials also live in old apartments with relatively poor WASH facilities compared to high-income elite communities. Large numbers of public latrines are also available in these neighborhoods. (We selected AGB officers' colony, Motijhil area, as a middle-income commercial community.) |
| Middle-income newly developed area | This is a newly-extended northern Thana (administrative sub district) and a planned square grid residential suburb of Dhaka which is geographically elevated from southern part of Dhaka. This newly-extended area has a relatively low population density, a good sewerage system and is less prone to flooding. (We selected the Uttarkhan area as the middle-income newly-extended community) |
| High-income community | For analysis, we combined high-income elite (Gulshan+ Dhanmondi), middle-income commercial/business community (Motijhil) and newly developed neighborhoods (Uttarkhan) together to define high-income communities. |
| Dhaka City Corporation [22] | In the Local Govt. (City Corporation) Amendment Act (2011), Dhaka City Corporation (DCC) was divided and re-created as Dhaka South City Corporation (DSCC) and Dhaka North City Corporation (DNCC) on 04.12.2011. |

**Table S2:** Definitions of different environmental samples collected for SaniPath deployment, 2017

| Pathway | Definitions |
| --- | --- |
| Municipal drinking water | Water supplied by the Water Supply and Sewerage Authority [WASA], including both “legal” and “illegal” connections. Water may be accessed through: piped water into compounds (including flexible pipes); public taps/standpoints (a formally designated water station in the community, provided by the government, or managed by someone in the community); or water vendors/trucks. |
| Non-municipal drinking water | Water that was not directly provided by WASA and was commonly used as drinking water in the neighborhood. These included shallow tube wells, submersible pump connected to deep tube wells, and 20 L commercially-available jars/bottles of water. |
| Bathing water | Water used for bathing by people in the community. This included WASA water and/or tubewell water commonly used as bathing water. Bathing water was stored or used directly from the source, as reported by the users during sample collection. |
| Surface water | Water collected from lakes and ponds |
| Drain water | Water from a channel carrying liquid and solid waste, including rainwater, floodwater, and sewage |
| Floodwater | Stagnant water that remains for at least one hour after raining |
| Communal /shared latrine | Communal latrines are accessed by any neighborhood resident. Shared latrines are accessed only by specific households. These latrines are not located within a household. |
| Raw produce | Vegetables that are commonly eaten without cooking. These vegetables do not have a shell or peel and grow above ground. Common produce items include cucumber, tomato, and coriander. |
| Street food [15,28] | Food sold on the street and commonly eaten by people in the community. Common street food we collected included *fuska* (a round puffed and fried crisp; a hole is created on the top to add a spiced sauce filling), *Chotpoti* (popular hot and sour snacks among the urban people in Bangladesh which is made up of potatoes, chickpeas, onions and chilies mixed with tamarind sauce), and *Jhalmuri* (mixture of puffed rice and a variety of spices, including peanuts, mustard oil, chili, onion, tomato, fresh ginger, salt, and/or lemon juice). |
| Soil | Soil/sand/mud was collected and analyzed from areas where people gather and/or where children play within the neighborhood. |

**Table S3:** Summary of information from the key informant interviews (KII) about the characteristics of the 10 study neighborhoods in Dhaka city, 2017.

| Characteristics | Latrine type | Open drain present | Do Children play in/near the drain | Common bathing water source | Common municipal drinking water source | Common non-municipal drinking water source | Common surface water source | Common types of produce eaten raw | Common types of street food eaten | Does it flood in this neighborhood? | Do children play in the flood-water? |
| --- | --- | --- | --- | --- | --- | --- | --- | --- | --- | --- | --- |
| Categories | Private latrine (1), Shared latrine (2), Public latrine (3) | Yes (1)<br>No (0) | Yes (1)<br>No (0) | WASA (1), Shallow Tubewell (2), Surface water (3) | WASA tap (1), WASA handpump (2), WASA Public tap (3) | Jar water (1), Deep borewell (2), Submersible pump (3), Bottled water (4) | Ditch (1), Pond (2), Lake (3), River (4), Canal (5) | Cucumber (1), tomato (2), coriander (3), carrot (4), green chili (5), Mint (6), Capsicum (7) | *Jhalmuri*(1), *Chotpoti*(2), *Fuska* (3), *Halim* (4), *Dal puri* (5), Pickle (6) | Yes (1)<br>No (0) | Yes (1)<br>No (0) |
| **Neighborhoods** |  |  |  |  |  |  |  |  |  |  |  |
| **Floating communities** |  |  |  |  |  |  |  |  |  |  |  |
| Gabtolli | 2, 3 | 1 | 0 | 1, 3 | 1 | 1 | 1 | 1, 3, 4, 5, 6 | 1, 2, 3, 4 | 1 | 1 |
| Kamalapur | 2, 3 | 1 | 1 | 1, 2 | Nil | 1, 2 | 1, 2, 5 | 1, 2, 3, 4, 5 | 1, 2, 3 | 1 | 1 |
| **Unstructured slums** |  |  |  |  |  |  |  |  |  |  |  |
| Kalshi | 2, 3 | 1 | 1 | 1, 2 | 1 | 1 | 3 | 1, 2, 4 | 1, 2, 3 | 1 | 1 |
| Shampur | 2 | 1 | 1 | 1, 2 | 1 | 1 | 2 | 1, 3, 4, 5 | 1, 3 | 1 | 1 |
| **Structured slums** |  |  |  |  |  |  |  |  |  |  |  |
| Badda | 1, 2, 3 | 1 | 1 | 1, 2 | 1, 2 | 1 | 1 | 1, 3, 4, 5 | 1, 2, 3 | 1 | 1 |
| Hazaribag | 1, 2 | 1 | 1 | 1, 2 | 1, 2, 3 | 1 | 4, 5 | 1, 2, 3, 4, 5 | 1, 2, 3, 5 | 1 | 1 |
| **Non-slum with poor WASH facilities** |  |  |  |  |  |  |  |  |  |  |  |
| Uttarkhan | 1, 2, 3 | 1 | 0 | 1, 2 | Nil | 3 | 2 | 1, 2, 3, 4, 5 | 1, 2, 3, 6 | 1 | 1 |
| Motijhil | 1, 2, 3 | 1 | 1 | 1 | 1, 3 | 1 | 2 | 1, 3, 4, 5 | 1, 2, 3, 4 | 1 | 1 |
| **Non-slum with improve WASH facilities** |  |  |  |  |  |  |  |  |  |  |  |
| Gulshan | 1, 2, 3 | 1 | 0 | 1 | 1 | 1, 4 | 3 | 1, 2, 4, 7 | 1, 2, 3 | 0 | 0 |
| Dhanmondi | 1, 2, 3 | 0 | 0 | 1, 3 | 1 | 1, 4 | 3 | 1, 2, 4, 7 | 1, 2, 3 | 1 | 1 |

*Jhalmuri*: mixture of puffed rice and a variety of spices, including peanuts, mustard oil, chili, onion, tomato, fresh ginger, salt and/or lemon juice
*Chotpoti*: popular hot and sour snacks among the urban people in Bangladesh, which is made of potatoes, chickpeas, onions, and chilies mixed with tamarind sauce
*Fuska*: a round, puffed and fried crisp; a hole is created on the top to add a spiced saucefilling
*Halim*: made of wheat, barley, meat (usually minced beef or mutton), lentils and spices, sometimes rice is also used.
*Dal puri*: an unleavened deep-fried bread along with lentils, onions and/or (rock) salt.

**Table S4:**
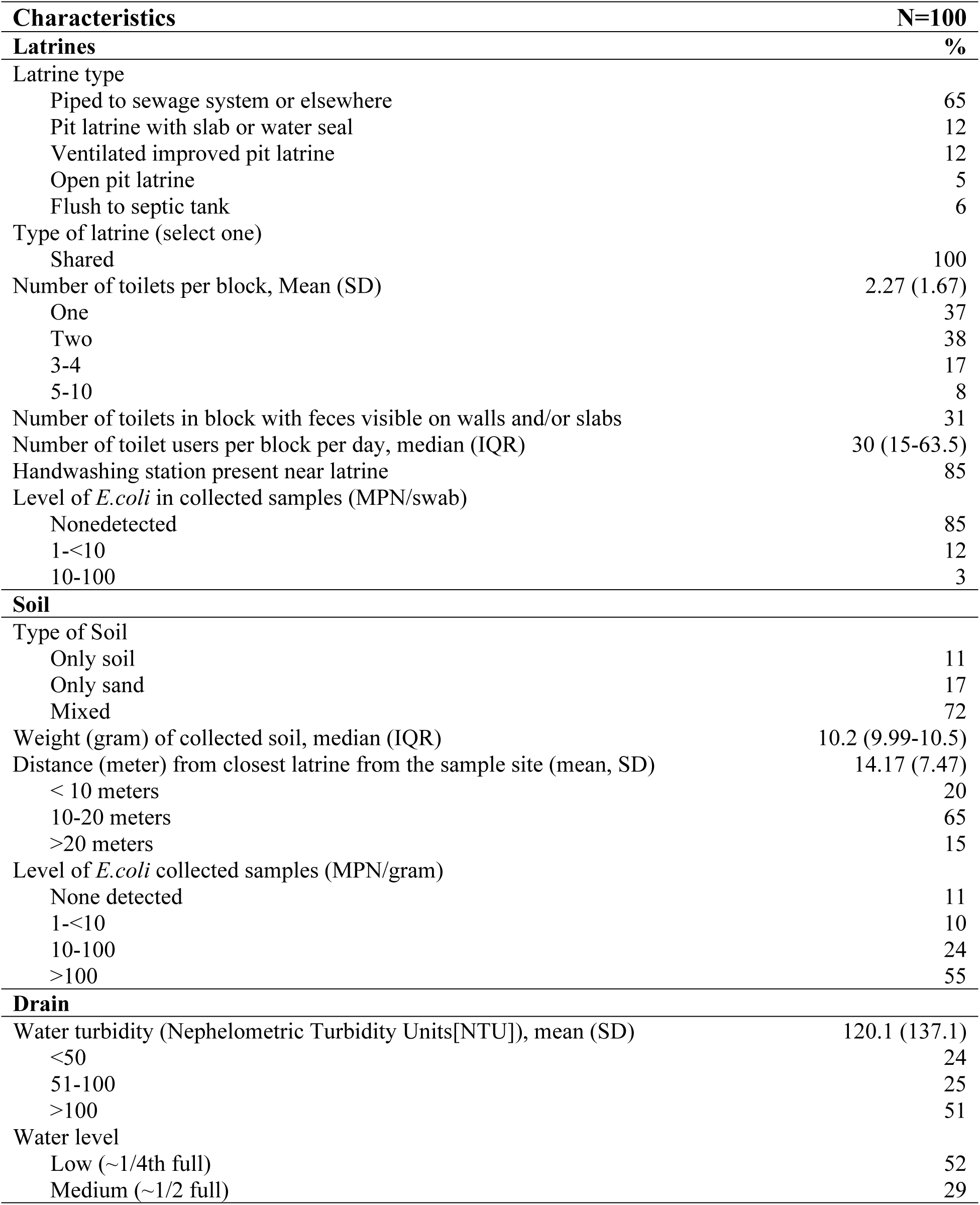
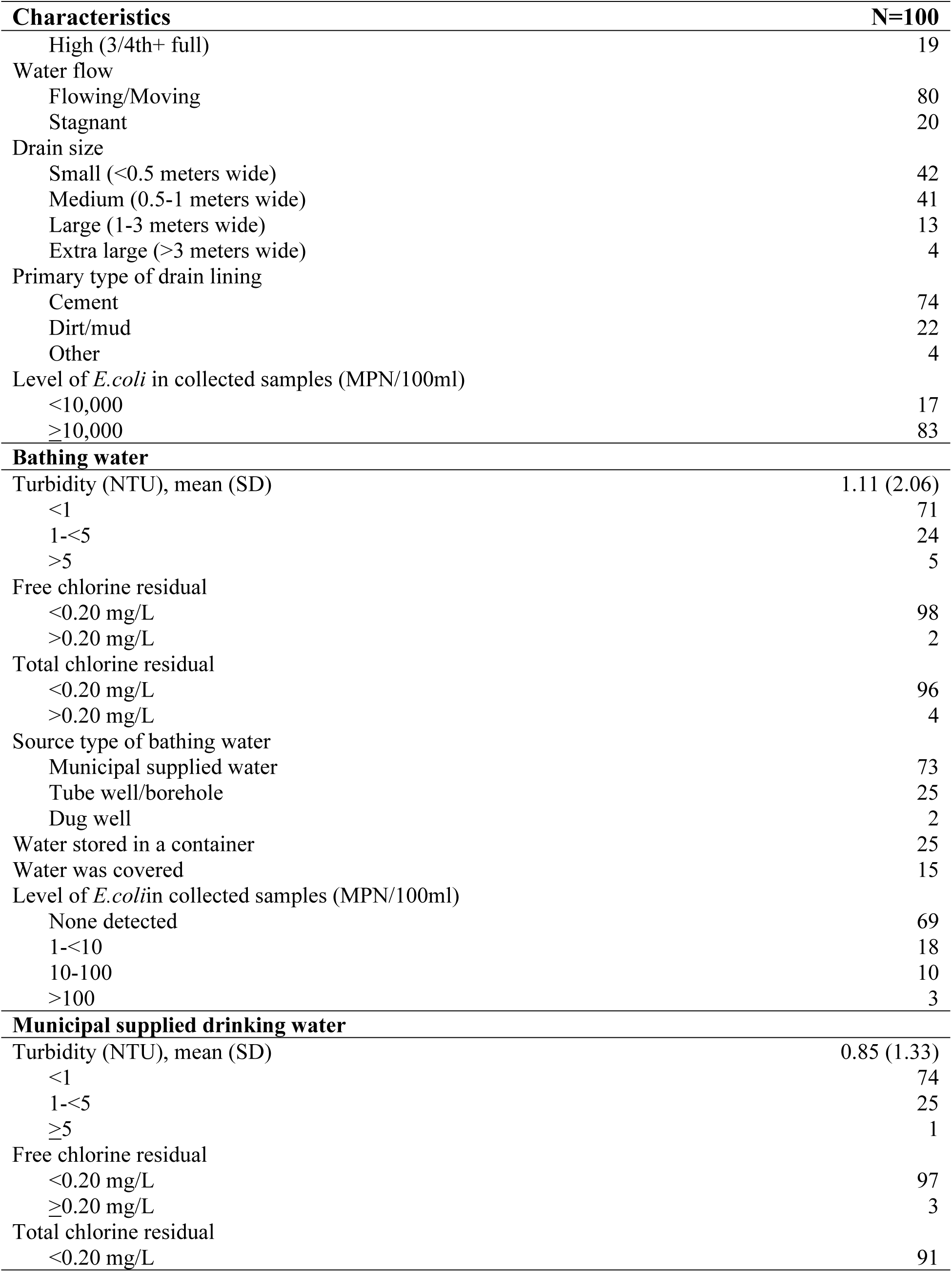
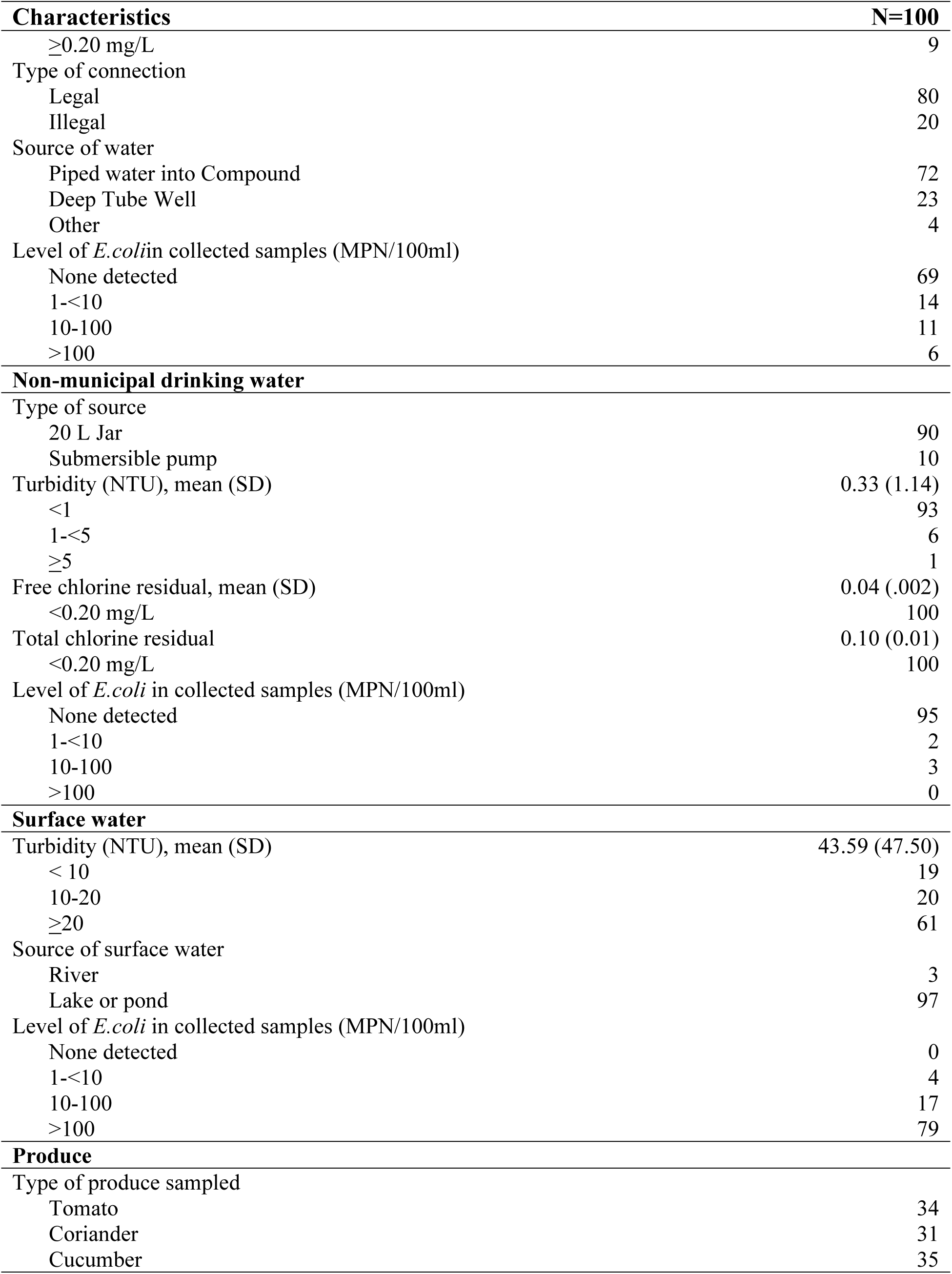
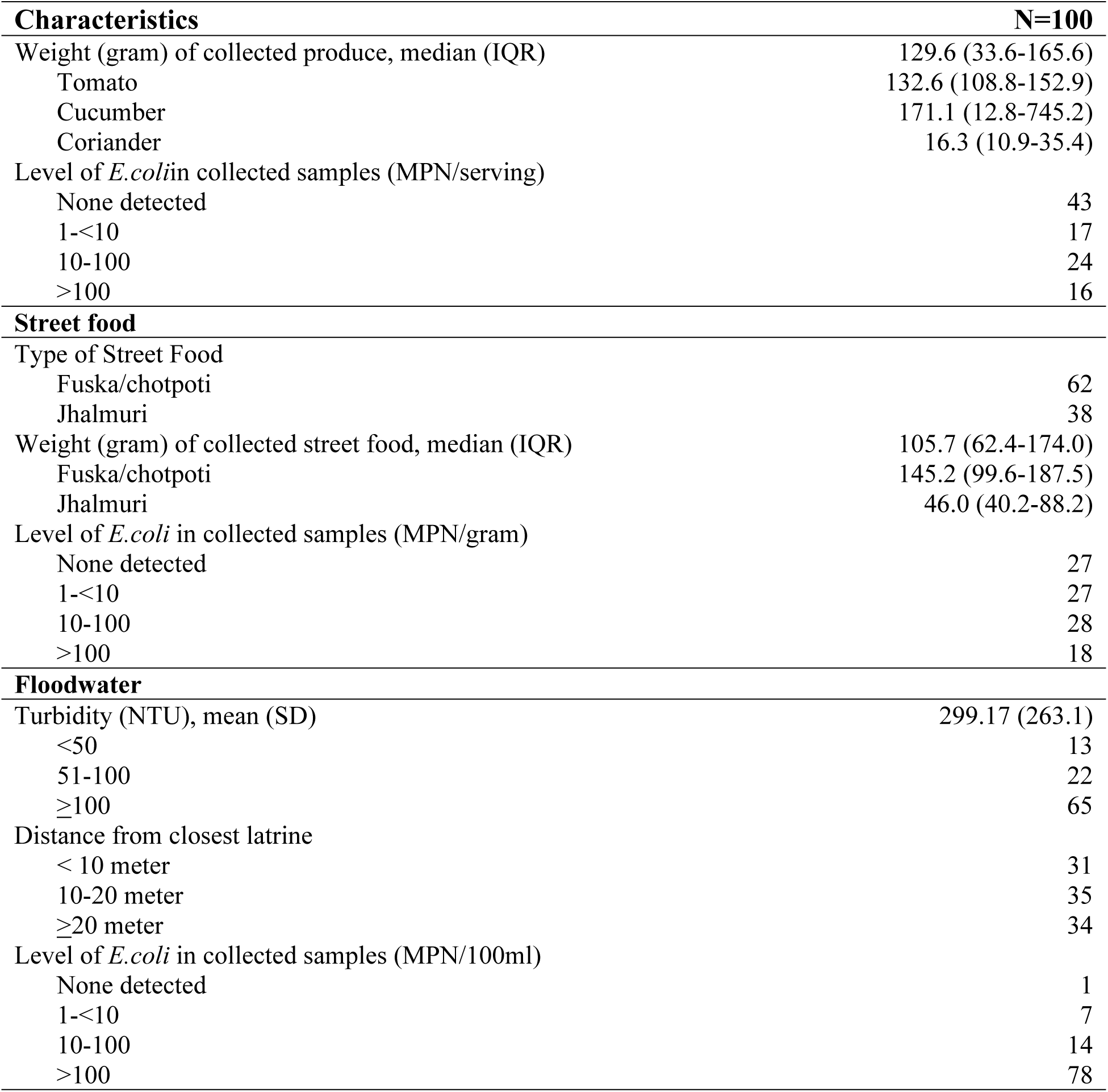
Summary of characteristics of the environmental samples in all 10 study neighborhoods of urban Dhaka, 2017.

| **Characteristics** | **N=100** |
| --- | --- |
| **Latrines** | **%** |
| Latrine type |  |
| Piped to sewage system or elsewhere | 65 |
| Pit latrine with slab or water seal | 12 |
| Ventilated improved pit latrine | 12 |
| Open pit latrine | 5 |
| Flush to septic tank | 6 |
| Type of latrine (select one) |  |
| Shared | 100 |
| Number of toilets per block, Mean (SD) | 2.27 (1.67) |
| One | 37 |
| Two | 38 |
| 3-4 | 17 |
| 5-10 | 8 |
| Number of toilets in block with feces visible on walls and/or slabs | 31 |
| Number of toilet users per block per day, median (IQR) | 30 (15-63.5) |
| Handwashing station present near latrine | 85 |
| Level of *E.coli* in collected samples (MPN/swab) |  |
| Nonedetected | 85 |
| 1-<10 | 12 |
| 10-100 | 3 |
| **Soil** |  |
| Type of Soil |  |
| Only soil | 11 |
| Only sand | 17 |
| Mixed | 72 |
| Weight (gram) of collected soil, median (IQR) | 10.2 (9.99-10.5) |
| Distance (meter) from closest latrine from the sample site (mean, SD) | 14.17 (7.47) |
| < 10 meters | 20 |
| 10-20 meters | 65 |
| >20 meters | 15 |
| Level of *E.coli* collected samples (MPN/gram) |  |
| None detected | 11 |
| 1-<10 | 10 |
| 10-100 | 24 |
| >100 | 55 |
| **Drain** |  |
| Water turbidity (Nephelometric Turbidity Units[NTU]), mean (SD) | 120.1 (137.1) |
| <50 | 24 |
| 51-100 | 25 |
| >100 | 51 |
| Water level |  |
| Low (~1/4th full) | 52 |
| Medium (~1/2 full) | 29 |
| High (3/4th+ full) | 19 |
| Water flow |  |
| Flowing/Moving | 80 |
| Stagnant | 20 |
| Drain size |  |
| Small (<0.5 meters wide) | 42 |
| Medium (0.5-1 meters wide) | 41 |
| Large (1-3 meters wide) | 13 |
| Extra large (>3 meters wide) | 4 |
| Primary type of drain lining |  |
| Cement | 74 |
| Dirt/mud | 22 |
| Other | 4 |
| Level of *E.coli* in collected samples (MPN/100ml) |  |
| <10,000 | 17 |
| ≥10,000 | 83 |
| **Bathing water** |  |
| Turbidity (NTU), mean (SD) | 1.11 (2.06) |
| <1 | 71 |
| 1-<5 | 24 |
| >5 | 5 |
| Free chlorine residual |  |
| <0.20 mg/L | 98 |
| >0.20 mg/L | 2 |
| Total chlorine residual |  |
| <0.20 mg/L | 96 |
| >0.20 mg/L | 4 |
| Source type of bathing water |  |
| Municipal supplied water | 73 |
| Tube well/borehole | 25 |
| Dug well | 2 |
| Water stored in a container | 25 |
| Water was covered | 15 |
| Level of *E.coli* in collected samples (MPN/100ml) |  |
| None detected | 69 |
| 1-<10 | 18 |
| 10-100 | 10 |
| >100 | 3 |
| **Municipal supplied drinking water** |  |
| Turbidity (NTU), mean (SD) | 0.85 (1.33) |
| <1 | 74 |
| 1-<5 | 25 |
| ≥5 | 1 |
| Free chlorine residual |  |
| <0.20 mg/L | 97 |
| ≥0.20 mg/L | 3 |
| Total chlorine residual |  |
| <0.20 mg/L | 91 |
| ≥0.20 mg/L | 9 |
| Type of connection |  |
| Legal | 80 |
| Illegal | 20 |
| Source of water |  |
| Piped water into Compound | 72 |
| Deep Tube Well | 23 |
| Other | 4 |
| Level of *E.coli* in collected samples (MPN/100ml) |  |
| None detected | 69 |
| 1-<10 | 14 |
| 10-100 | 11 |
| >100 | 6 |
| **Non-municipal drinking water** |  |
| Type of source |  |
| 20 L Jar | 90 |
| Submersible pump | 10 |
| Turbidity (NTU), mean (SD) | 0.33 (1.14) |
| <1 | 93 |
| 1-<5 | 6 |
| ≥5 | 1 |
| Free chlorine residual, mean (SD) | 0.04 (.002) |
| <0.20 mg/L | 100 |
| Total chlorine residual | 0.10 (0.01) |
| <0.20 mg/L | 100 |
| Level of *E.coli* in collected samples (MPN/100ml) |  |
| None detected | 95 |
| 1-<10 | 2 |
| 10-100 | 3 |
| >100 | 0 |
| **Surface water** |  |
| Turbidity (NTU), mean (SD) | 43.59 (47.50) |
| < 10 | 19 |
| 10-20 | 20 |
| ≥20 | 61 |
| Source of surface water |  |
| River | 3 |
| Lake or pond | 97 |
| Level of *E.coli* in collected samples (MPN/100ml) |  |
| None detected | 0 |
| 1-<10 | 4 |
| 10-100 | 17 |
| >100 | 79 |
| **Produce** |  |
| Type of produce sampled |  |
| Tomato | 34 |
| Coriander | 31 |
| Cucumber | 35 |
| Weight (gram) of collected produce, median (IQR) | 129.6 (33.6-165.6) |
| Tomato | 132.6 (108.8-152.9) |
| Cucumber | 171.1 (12.8-745.2) |
| Coriander | 16.3 (10.9-35.4) |
| Level of *E.coli* in collected samples (MPN/serving) |  |
| None detected | 43 |
| 1-<10 | 17 |
| 10-100 | 24 |
| >100 | 16 |
| **Street food** |  |
| Type of Street Food |  |
| Fuska/chotpoti | 62 |
| Jhalmuri | 38 |
| Weight (gram) of collected street food, median (IQR) | 105.7 (62.4-174.0) |
| Fuska/chotpoti | 145.2 (99.6-187.5) |
| Jhalmuri | 46.0 (40.2-88.2) |
| Level of *E.coli* in collected samples (MPN/gram) |  |
| None detected | 27 |
| 1-<10 | 27 |
| 10-100 | 28 |
| >100 | 18 |
| **Floodwater** |  |
| Turbidity (NTU), mean (SD) | 299.17 (263.1) |
| <50 | 13 |
| 51-100 | 22 |
| ≥100 | 65 |
| Distance from closest latrine |  |
| < 10 meter | 31 |
| 10-20 meter | 35 |
| ≥20 meter | 34 |
| Level of *E.coli* in collected samples (MPN/100ml) |  |
| None detected | 1 |
| 1-<10 | 7 |
| 10-100 | 14 |
| >100 | 78 |

**Table S5:**
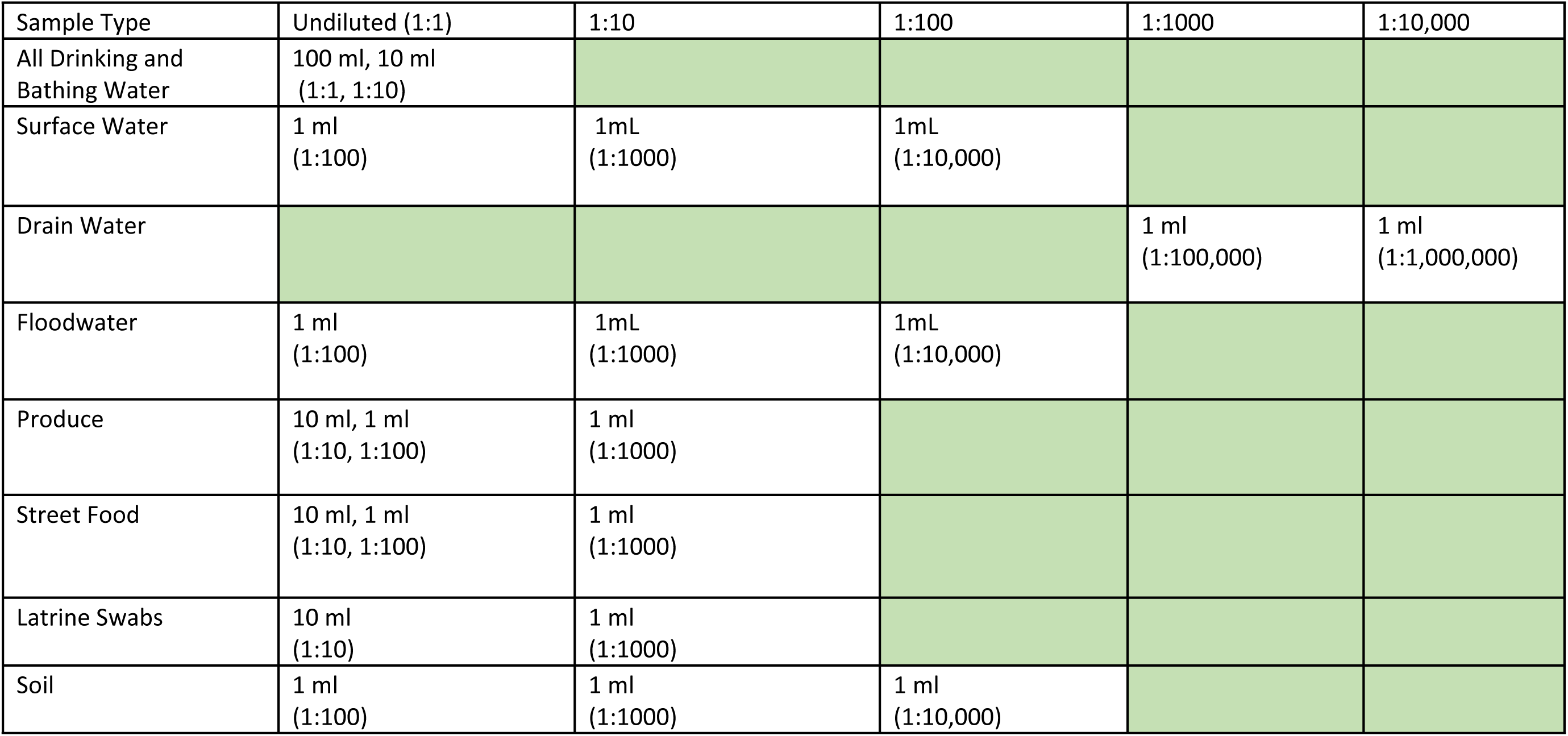
Dilutions tested of different types of environmental samples

**Figure S1:**
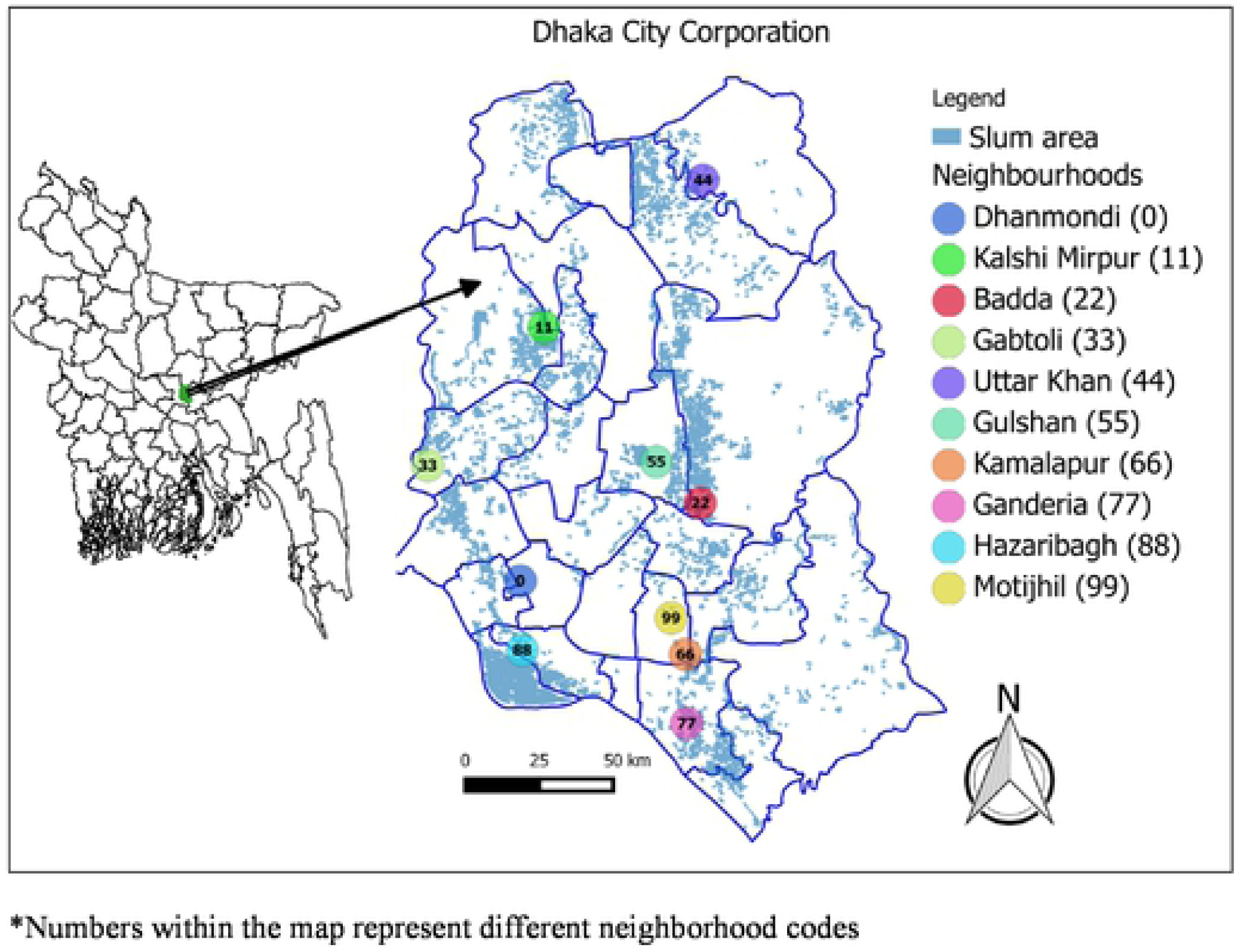
Locations of 10 study neighborhoods* in Dhaka city, 2017

